# Enantiomer-Specific Malathion Degradation by Gut Microbes of the Colorado Potato Beetle

**DOI:** 10.1101/2025.03.04.641399

**Authors:** Alison Blanton, Maria Olds, Adriana Martinez, Joshua Putman, Daniel Armstrong, Alison Ravenscraft

## Abstract

Symbiotic microbes play pivotal roles in insect ecology, including the detoxification of insecticides, which reduces target host mortality and diminishes the efficacy of chemical pest control agents. Quantifying the prevalence of symbiont-mediated insecticide detoxification across the microbiome is necessary to understand its contributions to pesticide resistance, and to understand how pesticides alter gut microbial communities. Here, we investigated the prevalence and mechanisms of pesticide degradation within the gut microbiota of the Colorado potato beetle (*Leptinotarsa decemlineata*), a significant agricultural pest that has driven ongoing insecticide innovation for decades. Beetles were collected from an organic farm in Tyler, TX, and 18 bacterial isolates representing the diversity of their gut microbiota were screened for their ability to degrade three common insecticides *in vitro*: imidacloprid, fenitrothion, and malathion. Among these, *Acinetobacter calcoaceticus, Pseudomonas protegens*, and an unnamed *Microbacterium* species degraded malathion as a sole carbon source, with distinct enantiomer-specific preferences. Untargeted GC-MS analysis revealed breakdown products, providing initial insights into the metabolic pathways utilized by these microbes. These findings suggest that microbial association with resistant insect hosts may select for microbial insecticide utilization, potentially enhancing resistance development in agricultural pests and influencing the surrounding soil microbiome.

## Introduction

The Colorado potato beetle (CPB) (*Leptinotarsa decemlineata*) is a major agricultural pest that primarily targets solanaceous crops, such as potatoes, eggplants, and tomatoes, often causing extensive economic damage (Alyokhin et al., 2013). This species first exhibited resistance to synthetic pesticides in 1952, when resistance to DDT was documented (Alyokhin et al., 2008). Since then, CPB populations have developed resistance to nearly all registered pesticides. Mechanisms of resistance include target-site mutations that reduce pesticide binding, enhanced enzymatic degradation via esterases, carboxylesterases, and monooxygenases, increased pesticide excretion, and cuticular modifications that limit chemical penetration (Alyokhin et al., 2008). The rapid evolution of pesticide resistance in the CPB is attributed to its high genetic diversity, fueled by rapidly evolving transposable elements, elevated genetic diversity, and a high gene expansion rate (Schoville et al., 2018). Furthermore, sublethal pesticide exposure under poor field conditions creates selective pressure that promotes epigenetic modifications conferring resistance, which are subsequently inherited by future generations (Chen et al., 2023). Additionally, the CPB’s exposure to plant toxins from its solanaceous diet may contribute to enhanced detoxification pathways, further compounding resistance. For instance, exposure to neonicotinoids has been linked to overexpression of cytochrome P450 genes, which are integral to xenobiotic metabolism (Zhu et al., 2016). In sum, a remarkable range of resistance and capability for rapid adaptation has positioned the CPB as a key driver in the development of modern insecticides.

A less explored but potentially significant contributor to CPB insecticide resistance is the role of its gut microbiota in pesticide degradation. Recent research has shown that the presence of *Lactococcus lactis* in the CPB gut correlates with reduced mortality following exposure to chlorpyrifos, and larvae inoculated with *L. lactis* exhibited similar resilience (Stara & Hubert, 2023). While the underlying mechanism remains unclear, it is hypothesized that gut microbes, including *L. lactis* and others, may directly degrade pesticides. The CPB gut microbiota encompasses bacteria such as *Lactococcus, Acinetobacter, Enterococcus*, and members of the *Enterobacteriaceae* family (Stara & Hubert, 2023), many of which have documented pesticide-degrading capabilities. For example, *Lactococcus lactis* has been shown to degrade organophosphorus pesticides (Pinto et al., 2019), and *Acinetobacter* can metabolize phenolic compounds, which are structural components of many pesticides (Argese et al., 2005; Hazai et al., 2004; Liu et al., 2016)

In this study, we quantified the prevalence of microbial pesticide detoxification across the gut microbiota of the CPB, focusing on three widely used insecticides: the organophosphates fenitrothion and malathion, and the neonicotinoid imidacloprid. Fenitrothion was selected because it represents the best understood case of microbially conferred pesticide resistance: *Caballeronia* bacteria evolved to degrade fenitrothion after exposure in agricultural soils, and when acquired by the bean bug (*Riptortus pedestris*), their natural host, they conferred resistance to the bug (Kikuchi et al., 2012). Malathion was chosen as a related organophosphate still in use in the United States (fenitrothion is no longer used). Finally, imidacloprid was included as a representative neonicotinoid with documented microbial degradation pathways.

It is important to note that microbes are not affected by insecticides in the same manner as host insects because microbes often lack the target sites that pesticides are designed to inhibit. Organophosphates, including malathion and fenitrothion, target the enzyme acetylcholinesterase (Kwong, 2002). Acetylcholinesterase aids in terminating nerve signals by binding to the neurotransmitter acetylcholine in postsynaptic nerve junctions (Bittner & Martyn, 2019). Organophosphates phosphorylate acetylcholinesterase irreversibly, preventing it from binding to acetylcholine and ultimately leading to the death of treated insects (Kwong, 2002). Similarly, neonicotinoid pesticides, including imidacloprid, bind to acetylcholine receptors in the synaptic cleft, also leading to nervous system dysregulation and death. Since microbes are single-celled organisms and do not have nervous systems, pesticides targeting the synapse should not adversely affect them. Instead, microbes can utilize pesticides as sources of carbon, nitrogen, or other nutrients.

Wild Colorado potato beetles (CPB) were collected from an organic farm in Tyler, TX, USA, to construct a library of gut bacterial isolates representative of the CPB microbiome’s diversity. We screened 18 isolates for degradation of fenitrothion, malathion, and imidacloprid. Additionally, the enantioselectivity of malathion-degrading isolates was tested, given the differential toxicity of malathion enantiomers. Bacterial growth curves were performed to determine whether these isolates could utilize the pesticides as sole carbon sources. To contextualize these findings, the composition of the CPB gut microbiota was characterized using Illumina amplicon sequencing, enabling an estimation of the relative abundance of pesticide-degrading genera in the wild beetle population.

## Methods

### Insect collection and bacterial isolation

Bacterial strains were isolated from the gut of Colorado potato beetles obtained from an organic farm in Tyler, Texas. Eleven individual beetles were surface sterilized with ethanol and homogenized under aseptic conditions. A small portion of the homogenate was serially diluted and plated on yeast-glucose (YG) agar; the remainder was immediately frozen at -80 C for culture-independent characterization of gut microbiome composition (described below).We picked at least one colony of each visible morphotype from each beetle and streaked it for isolation to obtain pure cultures. To preserve the isolates, we inoculated single colonies into YG broth, grew the cultures overnight, and froze the resulting bacterial suspension with a final concentration of 25% glycerol.

To identify the isolates, boiled cells were amplified with the universal bacterial 16S primers 327F (5’-ACACGGYCCARACTCCTAC-3’) and 936R (5’-TTGCWTCGAATTAAWCCAC-3’). The PCR recipe was 1 uL boiled cells, 0.4 µM forward primer, 0.4 µM reverse primer, 1 mM dNTPs, 1X New England Biolabs (NEB) buffer, and 1X NEB OneTaq in a final volume of 25 µL. The thermocycler settings were denaturation at 94 C for 1 min, 35 cycles of denaturation at 94 C for 30 s, annealing at 49 C for 30 s, and extension at 68 C for 1.5 min with a final extension at 68 C for 5 min. PCR products were sequenced by the Life Science Core Facility at the University of Texas at Arlington. Low-quality ends were trimmed, forward and reverse sequences were aligned, and the consensus sequences were searched against the NCBI database using BLAST. We chose 18 isolates to screen for pesticide degradation. These were selected with the goal of surveying the common members of the CPB microbiota. We avoided selecting multiple representatives of the same genus which were isolated from the same individual beetle, since these were likely to be duplicates of the same organism.

### Screening Media

To screen for insecticide degradation, we prepared an optimized version of M9 minimal media. The base media was composed of 3.3 g sodium phosphate dibasic (Na_2_HPO_4_), 1.5 g potassium phosphate monobasic (KH_2_PO_4_), 0.25 g sodium chloride (NaCl), and 0.50 g ammonium chloride (NH_4_Cl) dissolved in 500 mL of deionized water. For this screen, the media was supplemented with 0.50 g each of glutamic acid, histidine, tryptophan, and methionine. This mixture was autoclaved at 121°C for 15 minutes. After cooling, the media was further supplemented with vitamins including nicotinic acid, riboflavin, thiamine, and pantothenate at a final concentration of 0.2 μM. Additionally, 260 μL of 1 M magnesium sulfate (MgSO_4_), 14 μL of 1 M calcium chloride (CaCl_2_), and 3 mL of a 20% glucose solution were added.

Stock solutions of malathion, fenitrothion, and imidacloprid were prepared by dissolving 1 mg of each insecticide in 100 μL of acetonitrile (ACN). These stock solutions were used to achieve final media concentrations of 300 μM for malathion, 150 μM for fenitrothion, and 75 μM for imidacloprid to account for varying signal strength between insecticides. The resulting media were used in all insecticide degradation screening experiments.

### Cell Preparation and Incubation

Fresh bacterial cells were prepared by incubating each isolate in yeast glucose (YG) broth for 24 hours. Following incubation, cells were pelleted by centrifugation at ∼3,000 rpm, and the rich media was removed. The cell pellets were resuspended in screening media and used to inoculate experimental tubes at an initial optical density (OD) of 0.1. Uninoculated tubes containing insecticide-supplemented media served as abiotic controls. All tubes were incubated in the dark at 230 rpm for 48 hours, and four replicates were performed for each isolate and control.

### Post-Incubation Insecticide Extraction

After incubation, cells were pelleted by centrifugation at 12,000 rpm for 5 minutes, and the supernatant was collected. To enhance the polarity of the aqueous phase, 1.5 mL of phosphate-buffered saline (PBS; 137 mM NaCl, 2.7 mM KCl, and 10 mM phosphate, pH ∼7.4) was added to each sample. An internal standard, 31 μL of EPN/ACN [4mg/ml] (O-ethyl-O-p-nitrophenyl phenylphosphonothionate, final sample concentration 25 μM), was included to control for extraction efficiency. For the organic layer, 2.75 mL of 99.9% ethyl acetate was added to the sample, vortexed for 30 seconds, and the organic layer was collected. The ethyl acetate wash was repeated for a total of three times.

The pooled organic extracts were dried under nitrogen using a TurboVap™ system with a flow rate gradient (1.7–3.5 L/min) for 48 minutes. Dried samples were reconstituted in 100 μL of acetonitrile for high-performance liquid chromatography (HPLC) analysis.

### HPLC detection method

HPLC was performed on the 1100 Infinity II set from Agilent (Santa Clara, CA. USA) including a binary pump, mobile phase degasser, 96 vial sample injector, column thermostat, and diode array UV detector. A personal computer drove the chromatographic system and handled data with the OpenLab CDS Chemstation software (Agilent).

For the achiral separation, an Agilent InfinityLab Poroshell 120 SB-Aq column was used having a column length of 1 cm and 2.1 mm internal diameter, and filled with superficially porous 2.7 µm particles. Analysis conditions include a flow rate of 0.5 mL/min, column temperature of 45°C, and signal recording of 210 nm for malathion and 263 nm for imidacloprid, fenitrothion, and deltamethrin (internal standard). Mobile phases were A: 100% water with 0.1% formic acid and B: 100% acetonitrile with 0.1% formic acid, and the gradients used were 40%B, to 90%B, hold 1 min, then reequilibrate to 40%B (fenitrothion and malathion) and from 10% to 90%B, hold 1 min, then reequilibrate to 40%B (imidacloprid).

### Degradation Classification

Isolates were categorized based on their insecticide degradation capacity over the course of 48hrs. “Weak degraders” were defined as those eliminating between 20% and 30% of the pesticide. “Midrange degraders” eliminated between 30% and 80% of the pesticide, while “strong degraders” removed 80% or more of the pesticide.

### Total degradation and enantiomeric excess over time

To investigate total degradation and enantiomeric excess of malathion over time, we extended the methods used in the *in vitro* pesticide degradation screen. Bacterial cultures were grown for 60 hours in 5 mL volumes of supplemented minimal media, with an initial optical density (OD) of 0.1. Samples of 1 mL were collected at four time points: 15, 30, 40, and 60 hours. Each experimental condition was tested in eight replicates, alongside four no-culture controls to account for abiotic degradation. Overall growth was monitored to ensure and validate trends in degradation.

### Chiral Separations of Malathion

For the chiral separation, we used a Chiralpack IC-3 column packed with 3 µm fully porous particles, with column dimensions of 4.6 mm internal diameter and 25 cm length, provided by Daicel (Chiral Technologies, West Chester, PA, USA). Analysis conditions included a 2.0 mL/min flow rate, a column temperature of 45°C, signal recording of 210 and 263 nm, and a mobile phase of 90:10 hexanes:2-propanol.

Pesticide degradation was assessed using both achiral (total pesticide breakdown) and chiral (relative excess of malathion enantiomers) analyses. Enantiomeric excess (ee) was calculated using equation 1:

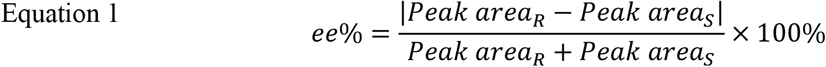

### In Vitro Carbon Utilization Assay

To assess the ability of malathion-degrading isolates to utilize malathion as a sole carbon source, we tested the three strongest degraders: *Acinetobacter calcoaceticus* (115f), *Pseudomonas protegens* (115d), and a *Microbacterium* species (103d). Each isolate was cultured overnight in yeast glucose (YG) broth, centrifuged and rinsed with respective treatment media, and diluted to an initial optical density (OD) of 0.2. The isolates were then monitored under four different conditions: YG broth as a rich media control, un-supplemented minimal media, minimal media supplemented with malathion and glucose as a secondary carbon source, and minimal media with malathion provided as the sole carbon source. The M9 minimal media formulation used in these experiments was consistent with the previously described screening media composition, except that glucose was omitted in the sole carbon source treatment.

### Evaluation of Metabolites produced by GCMS

To identify the metabolites produced by malathion-degrading isolates, we evaluated the breakdown of malathion by the three strongest degraders—*Acinetobacter calcoaceticus* (115f), *Pseudomonas protegens* (115d), and *Microbacterium* sp. (103d)—over the course of 96 hours. Each isolate was cultured overnight in yeast-glucose (YG) broth, centrifuged, rinsed with treatment-specific media, and diluted to an initial optical density (OD) of 0.1 in 6 mL of minimal media supplemented with malathion and glucose as a secondary carbon source. Media samples (1 mL) were extracted using the same extraction method and analyzed at the following time points: 20, 40, 50, 68, and 96 hours. Three controls were prepared alongside samples containing the bacterial isolate with malathion added: a no culture control (media only), a malathion-only control (no bacteria), and a bacteria-only control (no malathion). Preparation volumes, durations, and conditions were kept consistent between controls and samples. Both the overall malathion concentration and the presence of degradation metabolites were determined using gas chromatography-mass spectrometry (GC-MS), with compound identification performed via the NIST library.

### GC/MS Analysis of Malathion Metabolites

For the GC/MS analysis, an SLB-5 chromatographic column was used [(30 m x 250 mm x 1 um film thickness), (Supelco, Saint Louis, Missouri). The gas chromatograph utilized was a 6890N, coupled with a 5975 mass spectrometer (Agilent Technologies, Santa Clara, California). Helium carrier gas was used at a flow rate of 0.6 mL/min. 1 microliter of each sample was injected, with a split ratio of 20:1. Oven temperature programming was as follows: 40C, hold for 5 min, ramp at 10C/min to 260, hold 1 min (total run time = 28 min). For the mass spectrometer settings, a solvent delay of 2 min was used, and the mass scan range was 45-400 amu. Data processing was performed using MSD Chemstation D.02.00.275. Files were integrated using an automated threshold to determine which peaks were library searched. For the integration parameters, the settings included an initial threshold of 15.0, initial peak width of 0.100 (min), initial area reject of 1000, and shoulder detection was set to off. All peaks meeting this criteria for both controls and samples were manually library searched and reviewed using the NIST library (2005), and results tabulated.

To determine which metabolic products resulting from exposure to malathion, the controls were used to screen out peaks observed in the samples: Any peak observed in the sample and in the control was not reported as a metabolic product of malathion by the bacterial isolate.

### Characterization of the gut microbiome

DNA was extracted from 38 individual beetles (including the 11 individuals from which bacterial isolates were obtained) with the DNeasy Blood and Tissue Kit (Qiagen, Germantown, MD). Blanks were interspersed among the insect samples and were processed and identically to, and simultaneously with, the samples, including all downstream lab work and sequencing. We used the data from these blanks to identify and remove contaminants derived from the reagents or the laboratory environment (described below).

We amplified the V3-V4 hypervariable region of the 16S rRNA with universal bacterial primers 341F (5’-CCTACGGGNGGCWGCAG-3’) and 785R (5’-GACTACHVGGGTATCTAATCC-3’ (Klindworth et al., 2013). An overhang adapter was attached to the primers to enable the addition of sample indexes during a second amplification step (as per Illumina Inc., 2013). The PCR recipe was 1 uL DNA template, 0.5 uM forward primer, 0.5 uM reverse primer, and 1X Q5 master mix (New England Biolabs) in a final volume of 20 uL. The thermocycler settings were denaturation at 98 C for 1 min, 30 cycles of denaturation at 98 C for 10 s, annealing at 50 C for 20 s, and extension at 72 C for 30 s with a final extension at 72 C for 2 min. PCR products were cleaned with magnetic beads (Rohland & Reich, 2012).

The cleaned product was amplified in a second short PCR to attach a sample-specific pair of 8-nucleotide barcodes to the forward and reverse ends of the amplicons (Hamady et al., 2008). The barcodes enable sequences to be assigned to samples during data processing. The recipe for this second PCR was 2.5 uL PCR product, 0.5 uM of each barcoded primer, and 1X Q5 master mix in 20 uL. The thermocycler program was the same as above except that the annealing temperature was 55 C and only 8 amplification cycles were performed. The PCR products were cleaned with magnetic beads, quantified fluorometrically (Qubit DS DNA HS assay, Qiagen), and an equal mass of each sample was bidirectionally sequenced on one of three 600-cycle paired-end Illumina runs on a Mi-Seq platform at the University of Texas at Arlington’s Life Science Core Facility.

We removed priming sites and poor-quality bases from the ends of the sequences using the program cutadapt (Martin, 2011) Downstream processing was performed in R using the “DADA2” package (Callahan et al., 2016). Reads that had an expected error score greater than 2 or contained any unassigned bases (Ns) were discarded, and the remaining reads were truncated at the first instance of a quality score less than 2 using the filterAndTrim function. Forward and reverse reads were merged and bacterial sequence variants were inferred using the DADA2 algorithm, which uses the run-specific error rare, quality scores, and number of times each sequence is observed to infer true biological sequences, allowing analysis at the level of bacterial strains. Data from the three Illumina runs were then combined using the mergeSequenceTables function. We used the removeBimeraDenovo function to perform de novo chimera checking and removal. Bacterial taxonomy was assigned using the RDP classifier with the SILVA nr99 v138 database as the training set(Quast et al., 2012; Q. Wang et al., 2007) .

Next, we identified and removed contaminants. Since the expected minimum length of the bacterial amplicon is about 400-430 bp, we discarded sequences that were less than 398 base pairs in length or greater than 430 bp in length, as well as sequences identified as mitochondria or chloroplasts. To identify contaminants from the laboratory environment or reagents, we used the R package “decontam” with a threshold of 0.05 (Davis et al., 2018) to compare the prevalence of each sequence variant in the insect samples versus the extraction blanks. We removed 64 sequence variants identified as contaminants.

To control for per-sample differences in sequencing depth, all samples were rarefied to a depth of 13,263 reads. This rarefaction depth was chosen based on visual inspection of rarefaction curves, selecting the lowest depth at which a sample was sequenced and the curves had flattened out. Two samples were lost because they did not meet the rarefaction depth, resulting in a final dataset of 36 beetles, including 9 of the 11 individuals from which the screened isolates originated.

## Results

### Bacterial isolations

We obtained and successfully identified 42 bacterial isolates from field-collected Colorado potato beetles. Isolates belonged to, or were closely related to, the genera *Pantoea, Enterobacter, Microbacterium, Pseudomonas, Erwinia, Klebsiella, Lelliottia, Serratia, Acinetobacter, Plantibacter*, and *Rhizobium* (Table 1). Of these, we selected 18 isolates to screen for pesticide degradation ability. These isolates were selected to represent the diversity of the wild Colorado potato beetle gut microbiota from the study site as determined by culture-independent high-throughput sequencing (Fig. 4). The focal isolates included representatives of all the isolated genera except for *Plantibacter* and *Rhizobium*, which were never detected in the amplicon sequencing data, and *Klebsiella* because we included the closely related and more common *Lelliottia* instead. Overall, the screened isolates represented 64% of the wild CPB microbiome at the bacterial genus level (Table 1). We note that an additional 9% of the amplicon sequences belonged to *Spiroplasma*, which often lives inside insect cells as a facultative symbiont and is difficult to grow extracellularly in culture.

**Table 1.** Number of bacterial genera that were isolated versus number that were screened for pesticide degradation, and comparison of overall screen composition versus the abundance of these genera in wild Colorado potato beetles. Percentages of Illumina reads were calculated using rarefied data. Percentages in brackets were not included in the total percentage of Illumina reads because we did not screen representatives of these genera for pesticide degradation.

| Genus | N isolated | N screened | % isos screened | % Illumina reads |
| --- | --- | --- | --- | --- |
| *Pantoea* | 11 | 5 | 27.8% | 12.61% |
| *Enterobacter* | 9 | 3 | 16.7% | 36.31% |
| *Microbacterium* | 5 | 2 | 11.1% | 0.03% |
| *Pseudomonas* | 4 | 2 | 11.1% | 1.40% |
| *Erwinia* | 3 | 1 | 5.6% | 1.67% |
| *Klebsiella* | 2 | 0 | 0% | [0.82%] |
| *Lelliottia* | 1 | 1 | 5.6% | 5.30% |
| *Serratia* | 3 | 3 | 16.7% | 5.40% |
| *Acinetobacter* | 2 | 1 | 5.6% | 1.80% |
| *Plantibacter* | 1 | 0 | 0% | [0%] |
| *Rhizobium* | 1 | 0 | 0% | [0%] |
| **Totals** | **42** | **18** | **100%** | **64%** |

### Screening for Fenitrothion, Malathion, and Imidacloprid Degradation in the CPB Microbiome

The degradation of fenitrothion, malathion, and imidacloprid was assessed across a representative set of bacterial isolates from the Colorado Potato Beetle microbiome (Fig 1). For imidacloprid, two out of 18 isolates—*Enterobacter* RM109c and *Pseudomonas* RM115d— exhibited weak degradation, reducing the pesticide levels to approximately 70–79% of the original concentration. Fenitrothion degradation was slightly more pronounced, with five weak degraders (reducing levels to 75–77%) and two mid-range degraders (leaving 58–68% of the pesticide remaining). In contrast, malathion showed the highest degradation activity, with five weak degraders, two mid-range degraders, and three strong degraders. The top two isolates, *Acinetobacter calcoaceticus* (115f) and *Pseudomonas protegens* (115d), completely degraded malathion within 48 hours.

**Figure 1:**
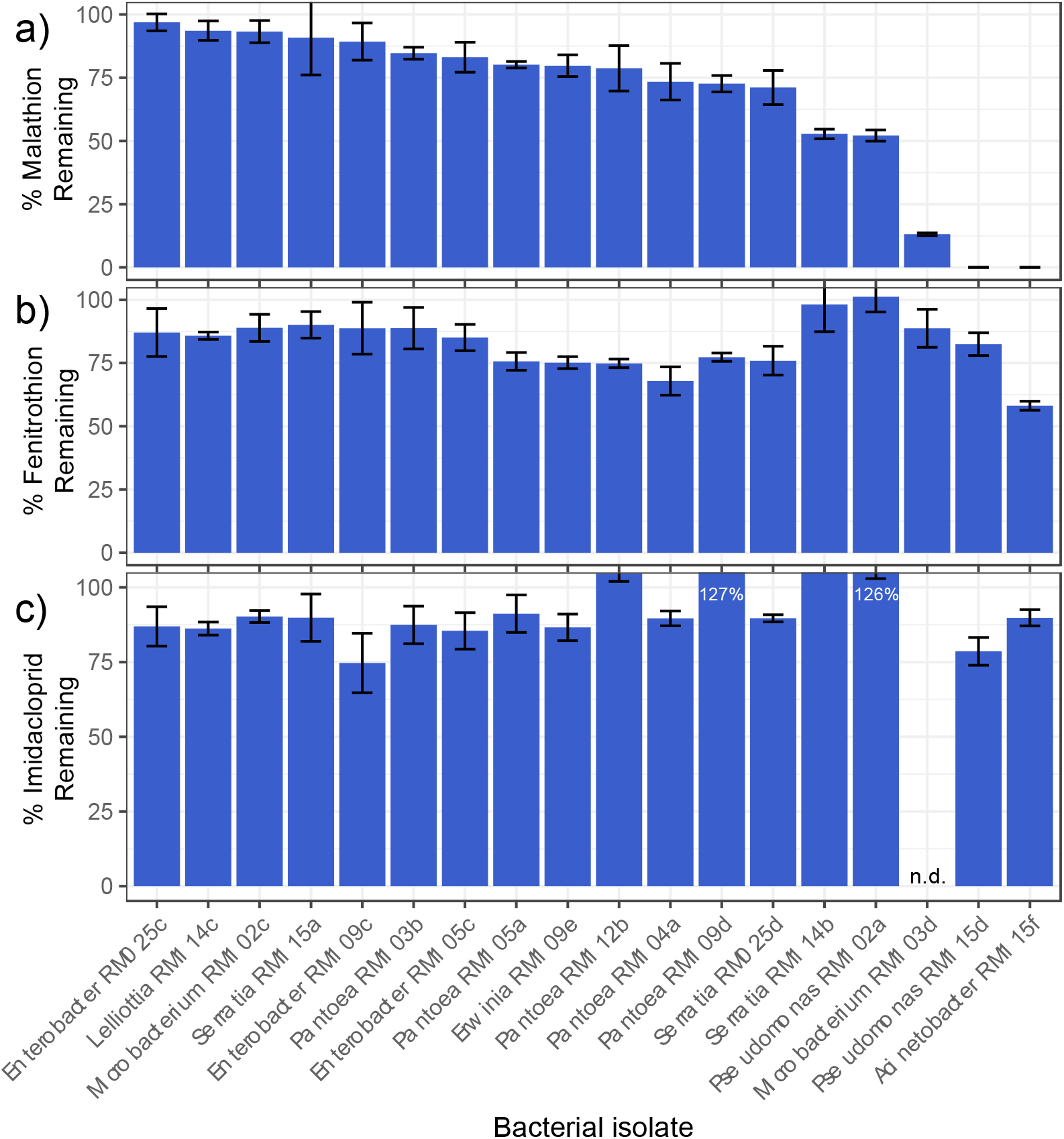
Degradation of imidacloprid (a), fenitrothion (b), and malathion (c) by bacterial isolates from the gut of Colorado potato beetles. Bars indicate the mean and error bars indicate the range of replicate measurements. n.d. = no data

### Evaluation of Degradation and Enantiomeric Excess Over Time

We assessed the total degradation of malathion and the enantiomeric preferences exhibited by the three strongest degrading isolates over 60 hours. Both *Pseudomonas protegens* (115d) and *Microbacterium* sp. (103d) maintained relatively steady degradation rates throughout the incubation period, while *Acinetobacter calcoaceticus* (115f) demonstrated delayed activity, with the majority of degradation occurring after 40 hours (Fig. 2a). The lack of complete degradation at 60hrs was attributed to slowed growth following the increase in culture volume from 1 mL to 6 mL to account for the media required to perform extra time points. Degradation trends at this volume remained consistent when compared to the 96-hour GC-MS results (Fig. 4).

**Figure 2:**
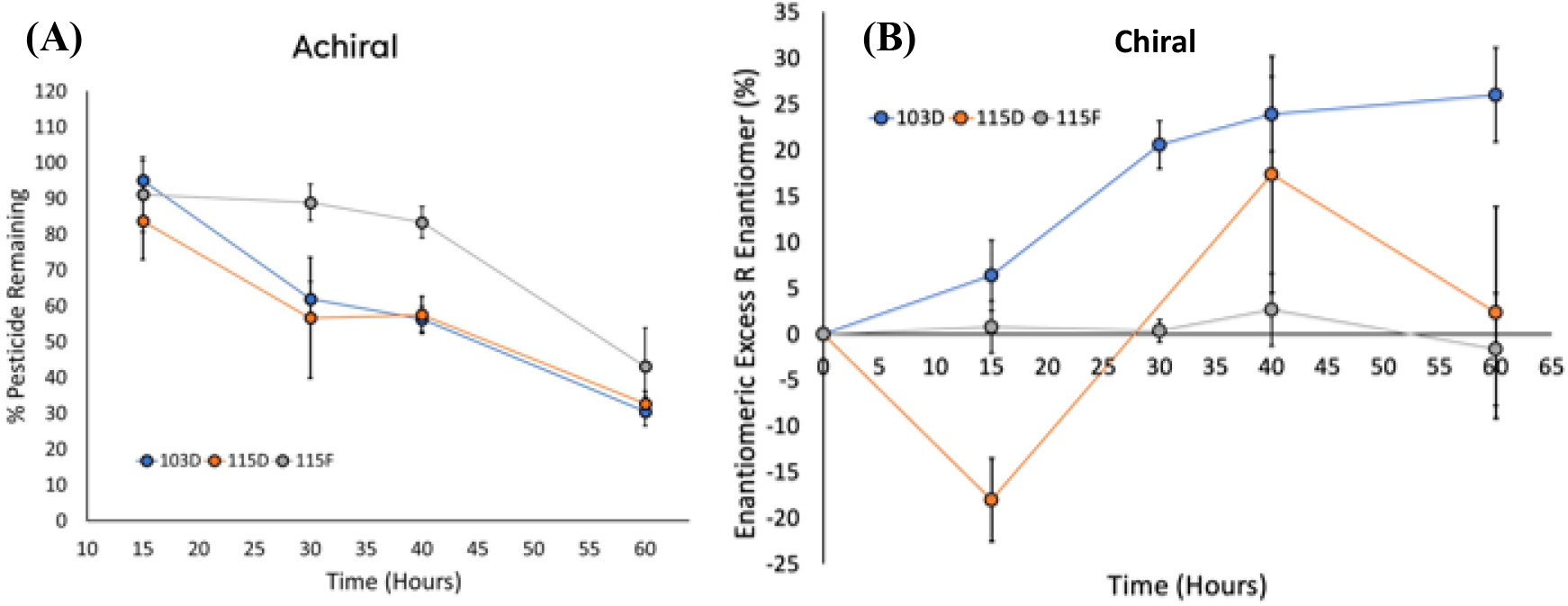
Time series of total malathion degradation and relative enantiomeric excess by isolates 115f (*Acinetobacter calcoaceticus)*, 115d (*Pseudomonas protegens*), and 103d (*Microbacterium sp*. Panel A: Total malathion concentration decreases over 60 hours for all three isolates. Panel B: Two isolates preferentially degraded one enantiomer of malathion, resulting in enantiomeric excess with error bars representing the relative standard deviation (RSD) between replicates. The enantiomeric excess (ee) was calculated using eqn. 1 (see methods)

Each isolate showed distinct preferences in enantiomer degradation. *Microbacterium* sp. (103d) preferentially degraded the less toxic S-enantiomer, leading to an increasing excess of the R-enantiomer over time (Fig. 2b). In contrast, *P. protegens* (115d) initially targeted the more toxic R-enantiomer, resulting in a transient excess of the S-enantiomer by 15 hours. This preference shifted to the S-enantiomer, becoming predominant until 40 hours, after which degradation returned to favoring the R-enantiomer. Finally, *A. calcoaceticus* (115f) exhibited no significant preference for either enantiomer.

### In vitro pesticide sole carbon source assay

All three isolates demonstrated growth under four tested conditions: rich media (yeast-glucose broth), minimal media, minimal media supplemented with both malathion and glucose as a secondary carbon source, and minimal media with malathion as the sole carbon source. Notably, all three were capable of utilizing malathion as their sole carbon source (Fig. S1). However, their growth dynamics varied significantly over time. *Acinetobacter calcoaceticus* (115f) exhibited delayed growth, remaining in lag phase until approximately 20 hours, after which it entered exponential growth and plateaued around 40 hours at a stable optical density (OD) of ∼0.9 through 48 hours. In contrast, both *Microbacterium* sp. (103d) and *Pseudomonas protegens* (115d) entered exponential growth within the first eight hours. *P. protegens* (115d) plateaued at an OD of ∼0.7 by 16 hours, maintaining this density through 48 hours. Meanwhile, *Microbacterium* sp. (103d) exhibited a more complex growth pattern, peaking at an OD of 0.28 around 20 hours before slightly declining to 0.26. Subsequently, it resumed exponential growth without plateauing, reaching an OD of 0.28 again at 48 hours. This secondary growth phase suggests potential diauxic growth driven by the consumption of malathion metabolites.

### Evaluation of Metabolites detected by GCMS

GC/MS analysis was used to identify and evaluate the occurrence of malathion metabolites produced by three bacterial isolates: *Acinetobacter calcoaceticus* (115f), *Pseudomonas protegens* (115d), and *Microbacterium* sp. (103d). Four breakdown products were detected across the isolates over the course of 96hrs (Table 2, Fig. 3): phosphorothioic acid, O,O,S-trimethyl ester (herafter “metabolite 1”, CAS 152-20-5), phosphorodithioic acid, O,O,S-trimethyl ester (“metabolite 2”, CAS 2953-29-9), butanedioic acid, mercapto-(“metabolite 3,” CAS 70-49-5), and butanedioic acid, [(dimethoxyphosphinothioyl)thio]-, 4-ethyl ester (“metabolite 4,” CAS 1190-29-0). For *Microbacterium* sp. (103d), metabolite 1 and metabolite 3 were detected after 20 hours of incubation, with metabolite 2 appearing at 35 hours, and all four metabolites detected by 52 hours (Fig. S2). In contrast, the only metabolite detected for *Pseudomonas protegens* (115d) at 20 hours was metabolite 4, which was observed at 20 hours and remained the sole detectable product until 96 hours, when metabolite 2 appeared (Fig. S2). Interestingly, *Acinetobacter calcoaceticus* (115f) exhibited no detectable metabolites until 96 hours, at which point metabolite 1, metabolite 2 and metabolite 4 were identified (Fig S2).

**Table 2.** Malathion metabolites identified by GC/MS for each isolate over a 96-hour incubation period. Xs indicate that a metabolite was detected.

| Metabolite | CAS ID | *Microbacterium* sp.<br>103d | *P. protegens*<br>115d | *A. calcoaceticus*<br>115F |
| --- | --- | --- | --- | --- |
| Phosphorothioic acid, O,O,S-trimethyl ester (metabolite 1) | 152-20-5 | X |  | X |
| Phosphorodithioic acid, O,O,S-trimethyl ester (metabolite 2) | 2953-29-9 | X | X | X |
| Butanedioic acid, mercapto- (metabolite 3) | 70-49-5 | X |  |  |
| Butanedioic acid, [(dimethoxyphosphinothioyl)thio]-, 4-ethyl ester (metabolite 4) | 1190-29-0 | X | X | X |

**Figure 3:**
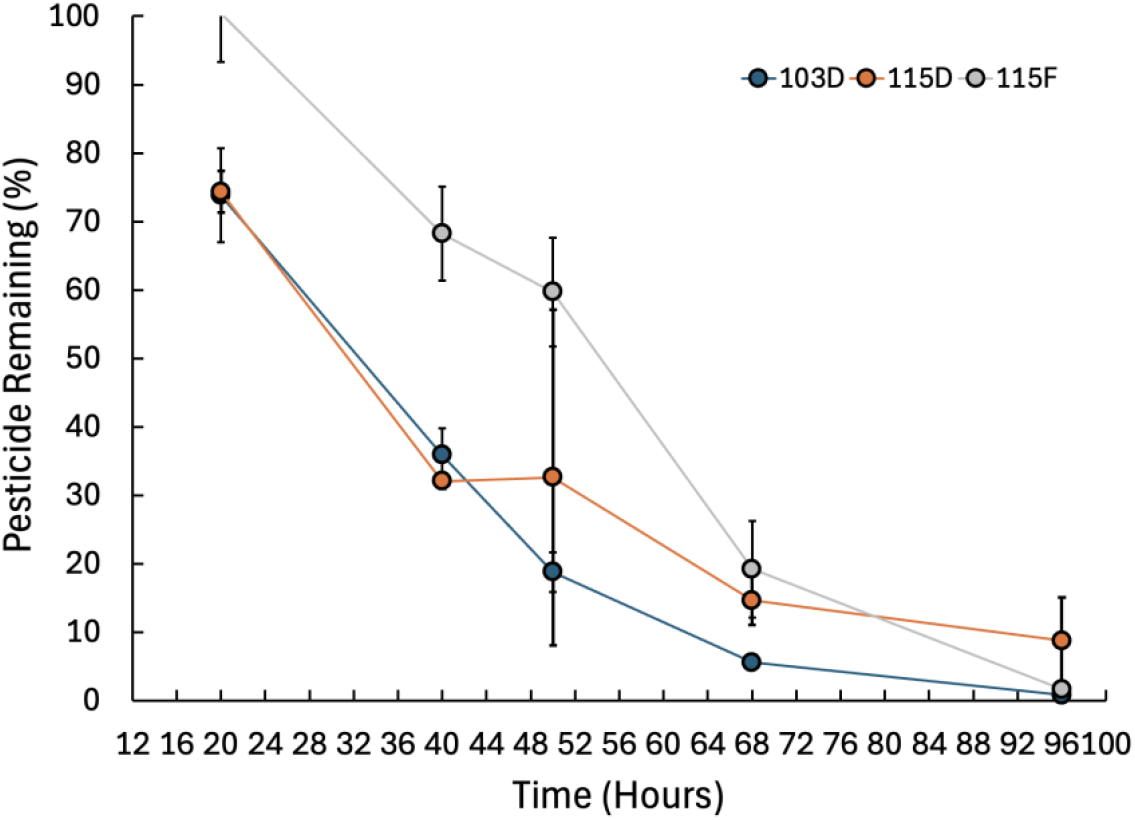
Tracking overall degradation of malathion using GC/MS during a 96-hour period to assess metabolites and confirm degradation trends record in smaller culture volumes.

**Figure 4.**
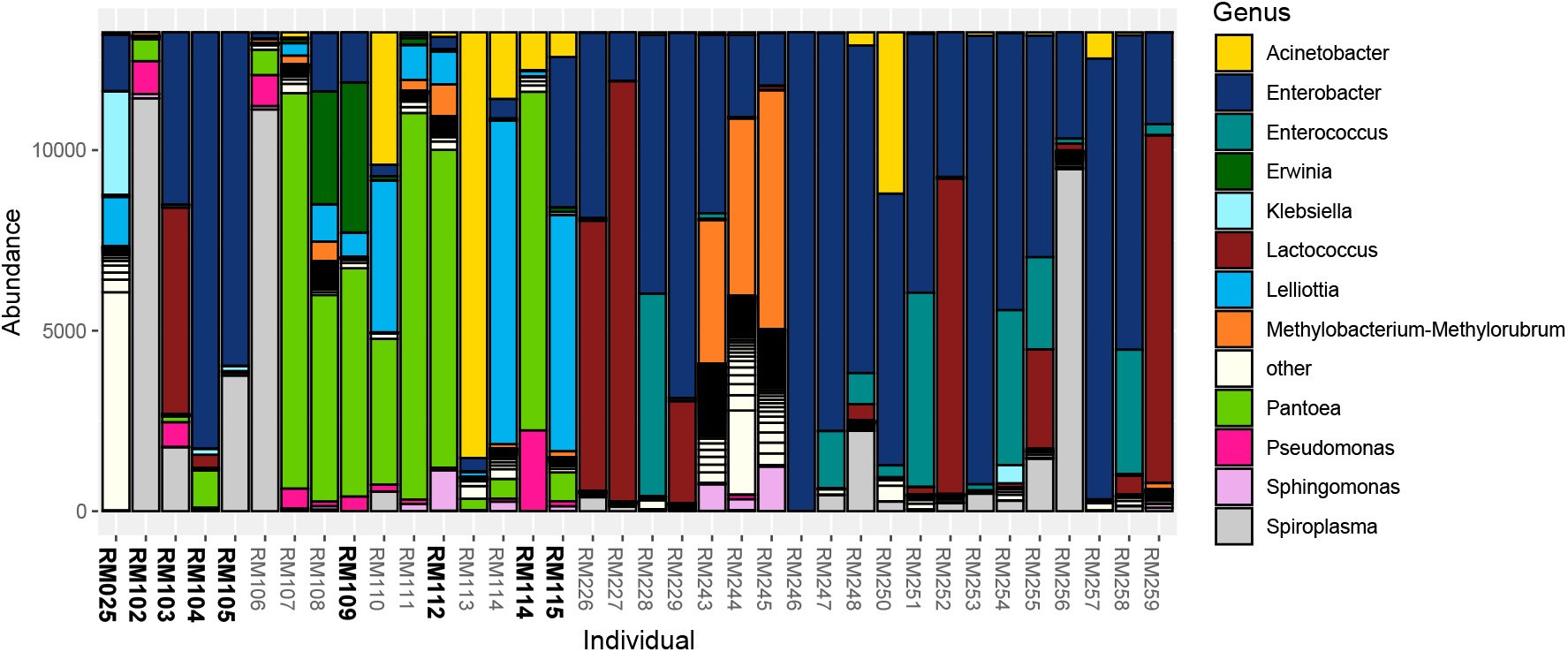
Relative abundances of bacterial genera in Colorado potato beetle at an organic farm in Tyler, TX. Each bar represents an individual insect. To control for differences in sequencing depth, all samples were rarefied to 13,263 reads. Colors indicate abundances of the most common bacterial genera (defined as those genera that accounted for at least 3% or more of the sequences in at least 2 beetles; rarer genera were assigned to the “other” category). Screened isolates were derived from the bolded individuals.

### Composition of the CPB gut microbiome

Overall, the most abundant bacterial genera in the Colorado potato beetle microbiome were *Enterobacter* (representing 36% of all DNA sequences after rarefaction), *Pantoea* (13%), *Lactococcus* (10%), *Spiroplasma* (9%), *Acinetobacter* (5%), *Lelliottia* (5%), *Enterococcus* (5%), *Methylobacterium* (4%), *Erwinia* (2%), *Pseudomonas* (1%) and *Serratia* (1%) (Fig 4). The isolates we tested in our pesticide degradation screen (Table 1) generally overlapped with these common genera, except for *Microbacterium* which was very uncommon (0.03%). We also note that *Spiroplasma* is often an endocellular symbiont, so the absence of *Spiroplasma* in our isolate library is expected.

## Discussion

Extensive prior research has shown that the CPB has evolved many innate mechanisms of resistance, but little is known about microbial contributions to resistance in this pest. To begin to address this question, we screened a representative set of gut bacterial isolates for the ability to degrade three insecticides. Most notably, three isolates identified as *Acinetobacter calcoaceticus, Pseudomonas protegens*, and *Microbacterium sp*. not only degraded most or all of the malathion in their cultures within 48 hours, but were able to utilize malathion as a sole carbon source. In addition, two of these isolates showed substantial and opposite preferential degradation of the malathion enantiomers, with *Pseudomonas protegens* (115d) preferentially breaking down the more toxic R-enantiomer and *Microbacterium* sp. preferentially metabolizing the less toxic S-enantiomer.

To our knowledge, one prior study has investigated conferred resistance to insecticides by the gut microbiome of CPB. It found that CPB better survived treatment with another organophosphate pesticide, chlorpyrifos, when they had a higher abundance of *Lactocaccus lactis* in their gut (Stara & Hubert, 2023). This suggests that *Lactocaccus lactis* might degrade chlorpyrifos, though it was not definitively demonstrated. In comparison, we identified three malathion degraders from genera that have not previously been implicated in pesticide degradation within the CPB (*Pseudomonas, Acinetobacter*, and *Microbacterium*). However, the first documented occurrence of organophosphorus degradation (OPD) genes was in an environmental *Pseudomonas*species (Yang et al., 2006), and other *Pseudomonas* species isolated from insects have been found to degrade diverse pesticides including organophosphates and pyrethroids (Itoh et al., 2018). Similarly, other *Acinetobacter* species have been found to degrade diverse pesticides in their environment including malathion (Sabit et al., 2011) neonicotinoids (M. Wang et al., 2011), pyrethroids and organochlorines (Gur Ozdal & Algur, 2022). All three genera are common environmental bacteria and warrant follow-up studies to test whether they can confer pesticide resistance to the Colorado potato beetle and other herbivorous insect pests.

We do not suggest that the CPB relies predominantly on its microbiome to detoxify pesticides. However, like many herbivorous insects, the CPB’s gut microbiota are facultative and are likely acquired from the beetle’s environment. Acquisition of gut colonists from local soil previously exposed to pesticides has been shown to confer fenitrothion resistance to the bean bug (*Riptortus pedestris*; (Kikuchi et al., 2012), and similar uptake of transient bacteria by the CPB could potentially provide occasional and/or supplementary detoxification to the beetle.

The preferential degradation of malathion by a subset of bacterial isolates may be attributed to at least two non-mutually exclusive factors. First, although the beetles were collected from an organic farm that does not use pesticides, they may have been previously exposed to malathion or structurally related insecticides from nearby conventional farms. Such exposure could have selected for microbiome-mediated degradation. In contrast, recent exposure to fenitrothion is unlikely, as its use has been banned in the United States since 2020. Second, malathion may inherently be more amenable to bacterial degradation due to short carbon tails on one side of the molecule.

Imidacloprid can be degraded by several microbial genera, including *Pseudomonas* and *Acinetobacter* (Pang et al., 2020). We were therefore surprised that we observed very little imidacloprid degradation. We speculate that this beetle population has not been recently exposed to imidacloprid or related neonicotinoids.

Regarding metabolite formation, malathion contains two ester bonds that are susceptible to hydrolysis by carboxylesterase enzymes, a common mechanism for pesticide breakdown. This enzymatic activity may partially explain the production of metabolites such as mercapto-butanedioic acid. In contrast, fenitrothion and imidacloprid lack ester bonds, which could account for their lower degradation rates if carboxylesterases are the primary enzymatic mechanism utilized by the bacterial isolates. Additionally, the modification of malathion’s phosphate ester group may involve phosphotriesterases, which are known to facilitate phosphate ester bond cleavage. This process could contribute to the production of phosphorothioic acid and phosphorodithioic acid, though achieving the final trimethyl moiety would likely require a secondary enzymatic interaction, such as methyltransferase activity. These enzymatic pathways have been well-documented in the microbial degradation of insecticides, particularly by soil bacteria (Ishag et al., 2016; Singh et al., 1999; Singh & Walker, 2006), which would align with the expected origin of the CPB microbiome. While not directly tested in this study, these proposed mechanisms represent a plausible route for gut microbial malathion metabolism.

In regard to the enantiomeric excess results, the degradation behavior of *P. protegens* in this study is particularly noteworthy. Due to the relatively short half-life of organophosphates compared to their predecessors, such as organochlorines, limited research has focused on the enantioselective degradation of chiral insecticides like malathion. Notably, the *R*-enantiomer of malathion has been reported to exhibit higher bioactivity and toxicity than the *S*-enantiomer, suggesting differential degradation rates may have toxicological implications in off-target organisms(Tong et al., 2023). Existing studies on malathion enantioselectivity have primarily examined degradation in soil and water microbial communities to infer broader environmental impacts(Ishag et al., 2016; Sun et al., 2012; F. Wang et al., 2023). One such study observed that the *S*-enantiomer degraded more rapidly than the *R*-enantiomer in both soil and water when exposed to a naturally occurring microbial community. However, the authors also noted that physiochemical factors, including pH and organic carbon content, significantly influenced the enantioselectivity of malathion degradation(Sun et al., 2012). These factors may partially account for the enantiomeric excess observed in our *in vitro* samples of *P. protegens* and *Microbacterium* sp., as bacteria can modulate their local pH within a certain range. However, pH modulation alone does not explain the switch in selective degradation observed in *P. protegens*, suggesting that additional microbial or enzymatic factors may contribute to this process influencing the initial preference for the R-enantiomer before switching to the S-enantiomer at 15hrs. However, the pattern we observed suggests that if these isolates confer pesticide resistance to the CPB, all else equal, isolate 103d (*Microbacterium* sp., which preferentially degrades the S-enantiomer) should be the least beneficial, 115f (*Acinetobacter calcoaceticus*, which exhibited no preference) should provide intermediate protection, and isolate 115d (*Pseudomonas protegens*, which preferentially degrades the R-enantiomer) should provide the most resistance to malathion.

Based on culture-independent microbiome sequencing, we estimate that the isolates we screened represent approximately 64% of the CPB microbiome at the genus level. Within this 64%, we found that 11% of microbial sequences belong to genera with members that mostly or completely degrade malathion, 24% belong to genera with members that moderately or weakly degrade malathion, and 33% belong to genera with members that weakly degrade fenitrothion. While these are back-of-the-envelope estimates with many caveats, they suggest that this beetle population’s gut bacteria have a high potential to degrade organophosphate insecticides. These high numbers are especially surprising given that we collected beetles on an organic farm that has not used synthetic pesticides for 15 years.

How representative might our findings be of pesticide degradation potential in other insect microbiota? Given its high degree of evolved pesticide resistance, CPBs in the field are likely treated with pesticides frequently, and at higher doses. Bacteria in the CPB gut might therefore be more likely to evolve the ability to break down insecticides as a nutrient source. However, the release of these microbes back into the environment (via insect secretions or death) could provide a source of bacterial isolates and genes that could be acquired by other insects (via ingestion) and other environmental bacteria (via horizontal gene transfer of pesticide degradation genes). Therefore, microbes residing in pesticide-resistant pests could potentially be an important contributor to the development of insecticide-degrading environmental bacteria and symbiont-mediated pesticide resistance.

## Conclusion

Colorado potato beetles (CPB) have developed resistance to over 50 pesticides (Chen et al., 2023). Much research has documented the beetle’s innate resistance mechanisms, but the frequency and magnitude of microbial contributions to the beetle’s resistance are unknown. We investigated the capacity of the Colorado potato beetle (*Leptinotarsa decemlineata*) gut microbiota to degrade two major families of insecticides that dominate the market today: organophosphates and neonicotinoids. Our findings reveal substantial malathion degradation, including complete degradation by two bacterial strains within 48 hours. Two of the top three degrader isolates exhibit enantioselective degradation, preferentially targeting different malathion enantiomers, but all three strains produce similar metabolites, suggesting the involvement of conserved enzymatic pathways These findings demonstrate the importance of investigating whether, and if so when and how much, gut bacteria contribute to pesticide resistance in the Colorado potato beetle and other herbivorous insect pests. Future research should assess whether the degradation patterns observed *in vitro* persist *in vivo* within insect hosts and in *in vitro* mixed cultures. Additionally, further studies should aim to identify the specific metabolic pathways utilized by these microbes to better understand their role in insecticide degradation and potential applications for pest management.

## Data Accessibility

Raw 16s Illumina amplicon sequences are available in the NCBI Sequence Read Archive under Project number PRJNA1225774. Bacterial 16S rRNA sequences are available on Genbank under accession numbers PV153584 and PV155211-PV155229.

## Acknowledgements

We thank Abiud Portillo for assistance with chemical methods development, Kaisy Martinez, Amelia Manns and Johnathan Adamson for assistance with insect collection and bacterial isolations, Spencer Reiling for help with pesticide detoxification screening, and Johnathan Adamson for assistance with Illumina library preparation. Support for this research was provided by a University of Texas at Arlington Maverick Bridge fellowship to Alison Blanton, a National Science Foundation CAREER grant to Alison Ravenscraft (2146512), a United States Department of Agriculture NIFA grant (2019-67013-29407) awarded to Martha S. Hunter, Alison Ravenscraft, and David Baltrus, and a USDA NIFA grant (2023-67013-39897) to Martha S. Hunter and Alison Ravenscraft.

## References

Alyokhin, A., Baker, M., Mota-Sanchez, D., Dively, G., Grafius, E., Alyokhin, A., Baker, M., Mota-Sanchez, D., Grafius, E., & Dively, G. (2008). Colorado Potato Beetle Resistance to Insecticides. J. Pot Res, 85, 395–413. 10.1007/s12230-008-9052-0

Alyokhin, Andrei., Vincent, Charles., & Giordanengo, Philippe. (2013). Insect pests of potato : global perspectives on biology and management. Elsevier /Academic Press.

Argese, E., Bettiol, C., Marchetto, D., De Vettori, S., Zambon, A., Miana, P., & Ghetti, P. F. (2005). Study on the toxicity of phenolic and phenoxy herbicides using the submitochondrial particle assay. Toxicology in Vitro, 19(8), 1035–1043. 10.1016/J.TIV.2005.05.004

Bittner, E. A., & Martyn, J. A. J. (2019). Neuromuscular Physiology and Pharmacology. Pharmacology and Physiology for Anesthesia: Foundations and Clinical Application, s412– 427. 10.1016/B978-0-323-48110-6.00021-1

Callahan, B. J., Mcmurdie, P. J., Rosen, M. J., Han, A. W., Johnson, A. J. A., & Holmes, S. P. (2016). dada2: high-resolution sample inference from illumina amplicon data. 13(7). 10.1038/nMeth.3869

Chen, Y. H., Cohen, Z. P., Bueno, E. M., Christensen, B. M., & Schoville, S. D. (2023). Rapid evolution of insecticide resistance in the Colorado potato beetle, Leptinotarsa decemlineata. In Current Opinion in Insect Science (Vol. 55). Elsevier Inc. 10.1016/j.cois.2022.101000

Davis, N. M., Proctor, Di. M., Holmes, S. P., Relman, D. A., & Callahan, B. J. (2018). Simple statistical identification and removal of contaminant sequences in marker-gene and metagenomics data. Microbiome, 6(1), 1–14. <10.1186/S40168-018-0605-2/FIGURES/6>

Gur Ozdal, O., & Algur, O. F. (2022). Biodegradation α-endosulfan and α-cypermethrin by Acinetobacter schindleri B7 isolated from the microflora of grasshopper (Poecilimon tauricola). 204, 159. 10.1007/s00203-022-02765-5

Hamady, M., Walker, J. J., Harris, J. K., Gold, N. J., & Knight, R. (2008). Error-correcting barcoded primers for pyrosequencing hundreds of samples in multiplex. Nature Methods 2008 5:3, 5(3), 235–237. 10.1038/nmeth.1184

Hazai, E., Gagne, P. V., & Kupfer, D. (2004). Glucuronidation of the oxidative cytochrome P450-mediated phenolic metabolites of the endocrine disruptor pesticide: methoxychlor by human hepatic UDP-glucuronosyl transferases. Drug Metabolism and Disposition: The Biological Fate of Chemicals, 32(7), 742–751. 10.1124/DMD.32.7.742

Ishag, A. E. S. A., Abdelbagi, A. O., Hammad, A. M. A., Elsheikh, E. A. E., Elsaid, O. E., Hur, J. H., & Laing, M. D. (2016). Biodegradation of Chlorpyrifos, Malathion, and Dimethoate by Three Strains of Bacteria Isolated from Pesticide-Polluted Soils in Sudan. Journal of Agricultural and Food Chemistry, 64(45), 8491–8498. 10.1021/ACS.JAFC.6B03334/SUPPL_FILE/JF6B03334_SI_001.PDF

Itoh, H., Hori, T., Sato, Y., Nagayama, A., Tago, K., Hayatsu, M., & Kikuchi, Y. (2018). Infection dynamics of insecticide-degrading symbionts from soil to insects in response to insecticide spraying. The ISME Journal 2018 12:3, 12(3), 909–920. 10.1038/s41396-017-0021-9

Kikuchi, Y., Hayatsu, M., Hosokawa, T., Nagayama, A., Tago, K., & Fukatsu, T. (2012). Symbiont-mediated insecticide resistance. Proceedings of the National Academy of Sciences of the United States of America, 109(22), 8618–8622. 10.1073/pnas.1200231109

Klindworth, A., Pruesse, E., Schweer, T., Rg Peplies, J., Quast, C., Horn, M., & Glö Ckner, F. O. (2013). Evaluation of general 16S ribosomal RNA gene PCR primers for classical and next-generation sequencing-based diversity studies. Academic.Oup.ComA Klindworth, E Pruesse, T Schweer, J Peplies, C Quast, M Horn, FO GlöcknerNucleic Acids Research, 2013•academic.Oup.Com, 41(1). 10.1093/nar/gks808

Kwong, T. C. (2002). Organophosphate pesticides: Biochemistry and clinical toxicology. Therapeutic Drug Monitoring, 24(1), 144–149. 10.1097/00007691-200202000-00022

Liu, Z., Xie, W., Li, D., Peng, Y., Li, Z., & Liu, S. (2016). Biodegradation of Phenol by Bacteria Strain Acinetobacter Calcoaceticus PA Isolated from Phenolic Wastewater. International Journal of Environmental Research and Public Health, 13(3), 300. 10.3390/IJERPH13030300

Martin, M. (2011). Cutadapt removes adapter sequences from high-throughput sequencing reads. EMBnet.Journal, 17(1), 10–12. https://journal.embnet.org/index.php/embnetjournal/article/view/200/479

Pang, S., Lin, Z., Zhang, Y., Zhang, W., Alansary, N., Mishra, S., Bhatt, P., & Chen, S. (2020). Insights into the Toxicity and Degradation Mechanisms of Imidacloprid Via Physicochemical and Microbial Approaches. Toxics 2020, Vol. 8, Page 65, 8(3), 65. 10.3390/TOXICS8030065

Pinto, G. D. A., Castro, I. M., Miguel, M. A. L., & Koblitz, M. G. B. (2019). Lactic acid bacteria - Promising technology for organophosphate degradation in food: A pilot study. LWT, 110, 353–359. 10.1016/J.LWT.2019.02.037

Quast, C., Pruesse, E., Yilmaz, P., Gerken, J., Schweer, T., Yarza, P., Rg Peplies, J., & Glö Ckner, F. O. (2012). The SILVA ribosomal RNA gene database project: improved data processing and web-based tools. Nucleic Acids Research, 41(D1), D590–D596. 10.1093/nar/gks1219

Rohland, N., & Reich, D. (2012). Cost-effective, high-throughput DNA sequencing libraries for multiplexed target capture. Genome Research, 22(5), 935–946. 10.1101/gr.128124.111

Sabit, H., Ahmed, O., Said, M., Shamseddin, A., Sabit, H. H., Said, O. A. M., Shamseldin, A. F., & Elsayed, K. (2011). Molecular identification of Acinetobacter isolated from Egyptian dumpsite as potential bacteria to degrade Malathion. International Journal of Academic Research, 3(4). https://www.researchgate.net/publication/265840587

Schoville, S. D., Chen, Y. H., Andersson, M. N., Benoit, J. B., Bhandari, A., Bowsher, J. H., Brevik, K., Cappelle, K., Chen, M. J. M., Childers, A. K., Childers, C., Christiaens, O., Clements, J., Didion, E. M., Elpidina, E. N., Engsontia, P., Friedrich, M., García-Robles, I., Gibbs, R. A., … Richards, S. (2018). A model species for agricultural pest genomics: the genome of the Colorado potato beetle, Leptinotarsa decemlineata (Coleoptera: Chrysomelidae). Scientific Reports 2018 8:1, 8(1), 1–18. 10.1038/s41598-018-20154-1

Singh, B. K., Kuhad, R. C., Singh, A., Lal, R., & Tripathi, K. K. (1999). Biochemical and molecular basis of pesticide degradation by microorganisms. Critical Reviews in Biotechnology, 19(3), 197–225. <10.1080/0738-859991229242/ASSET//CMS/ASSET/7F87C59C-8A4F-42FA-90C3>-32AB81607E69/0738-859991229242.FP.PNG

Singh, B. K., & Walker, A. (2006). Microbial degradation of organophosphorus compounds. FEMS Microbiology Reviews, 30(3), 428–471. 10.1111/J.1574-6976.2006.00018.X

Stara, J., & Hubert, J. (2023). Does Leptinotarsa decemlineata larval survival after pesticide treatment depend on microbiome composition? Pest Management Science, 79(12), 4921– 4930. 10.1002/PS.7694

Sun, M., Liu, D., Zhou, G., Li, J., Qiu, X., Zhou, Z., & Wang, P. (2012). Enantioselective degradation and chiral stability of malathion in environmental samples. Journal of Agricultural and Food Chemistry, 60(1), 372–379. 10.1021/JF203767D/ASSET/IMAGES/LARGE/JF-2011-03767D_0003.JPEG

Tong, Z., Shen, Y., Meng, D. D., Yi, X. T., Sun, M. N., Dong, X., Chu, Y., & Duan, J. S. (2023). Ecological threat caused by malathion and its chiral metabolite in a honey bee-rape system: Stereoselective exposure risk and the mechanism revealed by proteome. Science of The Total Environment, 874, 162585. 10.1016/J.SCITOTENV.2023.162585

Wang, F., Li, X., Jiang, S., Han, J., Wu, J., Yan, M., & Yao, Z. (2023). Enantioselective Behaviors of Chiral Pesticides and Enantiomeric Signatures in Foods and the Environment. Journal of Agricultural and Food Chemistry, 71(33), 12372–12389. 10.1021/ACS.JAFC.3C02564/ASSET/IMAGES/LARGE/JF3C02564_0004.J PEG

Wang, M., Yang, G., Wang, X., Yao, Y., Min, H., & Lu, Z. (2011). Nicotine degradation by two novel bacterial isolates of Acinetobacter sp. TW and Sphingomonas sp. TY and their responses in the presence of neonicotinoid insecticides. World Journal of Microbiology and Biotechnology, 27(7), 1633–1640. 10.1007/S11274-010-0617-Y/TABLES/2

Wang, Q., Garrity, G. M., Tiedje, J. M., & Cole, J. R. (2007). NaïveNaïve Bayesian Classifier for Rapid Assignment of rRNA Sequences into the New Bacterial Taxonomy †. APPLIED AND ENVIRONMENTAL MICROBIOLOGY, 73(16), 5261–5267. 10.1128/AEM.00062-07

Yang, C., Liu, N., Guo, X., & Qiao, C. (2006). Cloning of mpd gene from a chlorpyrifos-degrading bacterium and use of this strain in bioremediation of contaminated soil. FEMS Microbiology Letters, 265(1), 118–125. 10.1111/J.1574-6968.2006.00478.X

Zhu, F., Moural, T. W., Nelson, D. R., & Palli, S. R. (2016). A specialist herbivore pest adaptation to xenobiotics through up-regulation of multiple Cytochrome P450s. Scientific Reports 2016 6:1, 6(1), 1–10. 10.1038/srep20421

